# *Labyrinthula merlionensis* sp. nov.: a novel labyrinthulid infecting marine diatoms

**DOI:** 10.64898/2026.04.28.721384

**Authors:** Clarence Wei Hung Sim, Marie Walde, Hanna Strindberg, Avneet Kaur, Sophie Le Panse, Priscillia Gourvil, Jens Jahren, Daniel Vaulot, Adriana Lopes dos Santos

## Abstract

Labyrinthulomycetes are a class of fungus-like heterotrophic protists from the Stramenopiles lineage, recognized for their ecological role as decomposers and contributors to nutrient cycling. They colonize various substrates, from seaweed to terrestrial environments, utilizing ectoplasmic networks for nutrient absorption. This study characterized a novel *Labyrinthula* strain associated with the marine diatom *Biddulphia*. Phylogenetic analysis of the full-length 18S rRNA gene positioned this strain as a new species, *Labyrinthula merlionensis* sp. nov. Scanning electron and light microscopy observations revealed bi-flagellated zoospores and spindle-shaped vegetative cells with ectoplasmic networks. Time-series observations of the interactions between *L. merlionensis* and *Biddulphia* were categorised into different phases: establishment, infection, and aggregation. Scanning electron and confocal microscopy observations during the infection phase established the use of ectoplasmic nets to target the marginal ridge regions between diatoms, and the detection of labyrinthulid cells within diatom frustules. These findings enhance the understanding of the diversity, morphology, and ecological roles of Labyrinthulomycetes, particularly their intra- and extra-cellular interactions with diatom hosts.

## INTRODUCTION

Labyrinthulomycetes are a class of fungus-like heterotrophic protists belonging to the Stramenopiles lineage (Adl et al., 2019). The name Labyrinthulomycetes describes one of the characteristic features present in some genera: a network of fine cytoplasmic threads extending from the cell bodies into the environment, resembling a labyrinth (Porter, 1990). This "slime net" originates from a unique structure known as the bothrosome or sagenogenetosome, and its function varies between organisms.

Currently, five orders are recognized within the class Labyrinthulomycetes: Labyrinthulida, Oblongichytriida, Thraustochytrida, Amphifilida, and Amphitremida (Adl et al., 2019). The first four orders are characterized by the presence of the bothrosome (Bennett et al., 2017). Only Labyrinthulida has evolved the ability to glide along ectoplasmic nets for motility (Tsui et al., 2009).

Many Labyrinthulomycetes feed saprotrophically and are consequently recognized for their contribution to nutrient cycling (Raghukumar and Damare, 2011). Their unique ectoplasmic network allows them to expand surface contact with the substrate, enhancing their efficiency in absorbing nutrients from a wide range of organic sources. Additional feeding modes include bacterivory, with some Labyrinthulomycetes such as *Thraustochytrium striatum* and *Aurantiochytrium mangrovei* being able to produce a transient phagogrophic ameboid stage (Tsui et al., 2009).

Labyrinthulomycetes are considered to have a cosmopolitan distribution (Pan et al., 2017). While originally viewed exclusively as marine, they have since been found in a range of habitats, including estuarine, freshwater and terrestrial ones (Douhan et al., 2009). They are found as free living cells or in association with organic detritus or organisms such as macroalgae, microalgae, seagrasses and even terrestrial plants.

The position of *Labyrinthula* on the symbiotic spectrum (Rueckert et al., 2019) remains uncertain. Few species-host relationships involving members of the order Labyrinthulida have been well characterized. The species *Labyrinthula zostera* is recognized as the causative agent of the seagrass wasting disease events of the 1930s (Martin et al., 2016; Muehlstein et al., 1991). Another pathogenic species, *L. terrestris*, was discovered in association with dying turfgrass in a golf course in southern California (Bigelow et al., 2005). Cells of *Labyrinthula* have been isolated from tissues of healthy macroalgae, turfgrass and seagrasses, which suggests a possible commensal or mutual relationship with their hosts (Bockelmann et al., 2012; Muehlstein et al., 1991). What triggers the opportunistic pathogenicity is uncertain: it could be related to stress and degraded health in the host (Sullivan et al., 2013). Another hypothesis is linked to specific lineages being more virulent than others, as pathogenicity assays and phylogenetic analyses provided evidence of a distinct pathogenic clade of *Labyrinthula* (Martin et al., 2016).

Microscopic observations of cell-to-cell interactions between *Labyrinthula* and diatoms have also been documented for over a century (Jepps, 1931). However, the nature of this interaction remains uncertain. Currently, *L. magnifica* (1975) and *L. diatomea* (2020) are the two only formally described species of *Labyrinthula* associated with diatoms. Although a molecular characterization including reference sequences for *L. diatomea* has been obtained (Popova et al., 2020), representative strains of *L. diatomea* and *L. magnifica* associated with their original hosts are not currently available, limiting our ability to study this interaction under controlled laboratory conditions.

Given the ecological role of diatoms in global primary production (Field et al., 1998; Smetacek, 1999; Tréguer et al., 2018), understanding the nature of their interaction with *Labyrinthula* is important. In this paper, we characterize morphologically and molecularly a stable co-culture system involving a new species of *Labyrinthula* — *L. merlionensis* sp. nov. — and its original diatom host *Biddulphia* sp. We further investigate the dynamics of their interaction using confocal microscopy.

## MATERIALS AND METHODS

### Sampling, isolation and establishment of *Labyrinthula*-diatom co-culture

The *Labyrinthula*-diatom co-culture was isolated from natural seawater off the west coast of Saint John’s Island in the Singapore Straits (1.217085^◦^N, 103.845365^◦^E) on 26 February 2020. Seawater surface temperature and conductivity were measured using a SonTek CastAway™CTD. A vertical profile over 10 m depth was obtained by taking measurements at a frequency of 5 Hz. Salinity (PSS-78) was calculated from temperature, conductivity, and pressure measurements using EOS-80 equations following manufacturer software. Mean temperature and salinity were depth-integrated by computing the average of the vertical profile (Martin et al., 2022).

Seawater was collected using a 5 L Niskin bottle at 1 m depth and transported in a 1 L pre-washed (10% v/v HCl followed by three rinses using MilliQ Ultrapure water) polypropylene bottle to the laboratory within 6 h. *Labyrinthula* cells were observed to adhere to diatom cells using an inverted light microscope. The aggregate of *Labyrinthula* and diatom cells were isolated using a pre-pulled glass micropipette. The aggregate was incubated in an untreated vented culture flask using L1 + silicate seawater medium (Guillard and Hargraves, 1993) and regularly checked for growth of non-targetted taxa. Once established, the co-culture was maintained at 22^◦^C 12h:12h light-dark cycles, renewed every 4 to 5 weeks. The co-culture was deposited in the Roscoff Culture Collection as RCC7798. The diatom was subsequently isolated from the co-culture using serial dilution and maintained under the same conditions.

### DNA extraction and long-read rRNA operon sequencing

*Labyrinthula* and diatom cells were harvested at varying ages of the co-culture (Table S1) to ensure sufficient and good quality DNA for diatom (during diatom exponential growth phase) and *Labyrinthula* (after diatom population decline). Cells were concentrated by centrifugation. Total nucleic acids were extracted using the Nucleospin Plant II kit (Macherey-Nagel, Düren, DE) following the manufacturer’s instructions. Extracted genomic DNA concentration was measured using PicoGreen™(Thermo Fisher Scientific, Waltham, MA, USA) to ensure sufficient genomic content (>10 ng *µ*L *^−^*^1^) required by the sequencing facility (Integrated Microbiome Resource, Dalhousie University, Canada). The extracted genomic DNA was kept at −80^◦^C and shipped in dry ice to the sequencing facility. A PCR to obtain the nearly full length nuclear 18S rRNA gene amplicon was done using barcoded forward primer NSF4/18 (5’-CTGGTTGATYCTGCCAGT-’3) and reverse primer EukR (5’- TGATCCTTCTGCAGGTTCACCTAC-’3) following the protocol used by the sequencing facility

(Comeau and Filloramo, 2023). Library preparation was done using Pacific Biosciences (PacBio) SMRTbell libraries with the Express TPK2.0. Circular concensus sequence (CCS) reads were generated from raw sequences from PacBio Sequel 2 using the standard software tools provided by the manufacturer.

CCS reads were processed with scripts written in the R language (R Core Team, 2020) using the dada2 package adapted for long read amplicon sequencing (Callahan et al., 2019). Primers were first removed from reads using removePrimers function with default parameters, including from reversecomplemented reads using orient = TRUE. Reads were filtered and trimmed using the filterAndTrim function with the following parameters: minQ = 3, minLen = 1000, maxLen = 3000, maxN = 0, rm.phix = FALSE, maxEE = 2. Sequencing error rates were learnt using learnErrors function with the following parameters: errorEstimationFunction = PacBioErrfun, qualityType = FastqQuality. Chimeric sequences were removed with the removeBimeraDenovo function using default parameters. The majority of the CCS reads were retained (Table S1). ASVs were taxonomically assigned using assignTaxonomy function with PR^2^ database version 5.0.1 (https://pr2-database.org, Guillou et al., 2012) as a reference. ASVs with low (*<* 80%) bootstrap support from the assignTaxonomy function at a given taxonomic level were reclassified to the next higher taxonomic level until the bootstrap value was ≥ 80%. To account for the generation of false and low abundance ASVs from the oversequencing of a co-culture, ASVs were clustered at 99% using the –cluster_size option of vsearch version 2.18.0 (Table S2). For *Labyrinthula* sp., the centroid ASV with no gaps in the 18S sequence and that had the highest number of ASV clustered together was used for taxa description and phylogeny (Figure S1). The table of ASVs, processed sequence, and their DNA read abundance across samples are also available in the same repository. R and shell scripts used for the processing of pacbio ASVs are available at https://github.com/clarencesimple/SIM_Labyrinthula_merlionensis/.

### Molecular characterization

Phylogenetic analysis was performed using the 18S rRNA operon. We retrieved 270 18S rRNA Gen-Bank sequences that were annotated as *Labyrinthula* in PR^2^ v5.1 (Guillou et al., 2012) based on the Labyrinthulomycetes phylogeny by Pan et al. (2017). Sample type (culture, isolate or environmental) and sample substrate (e.g. water column, sediment, seagrass) of these sequences were manually curated. Sequences longer than 900 bp were selected and duplicated sequences were removed. Sequences found to be chimeric through preliminary alignment with MAFFT using default settings (Katoh and Standley, 2013) were also removed. This was done by visual inspection of high mismatches with the common consensus and verifying by BLASTing the sequence, to check if the sequence has any confident matches (>90% identity) with Labyrinthulomycetes. The *Labyrinthula* 18S sequence from co-culture of this study was also included. The 120 sequences were clustered at 99% identity using the –cluster_fast option of vsearch version 2.18.0, resulting in 41 clusters (centroid ASVs). Sequences that did not form any cluster and formed an isolated clade in preliminary iterations of the phylogenetic tree were removed. Sequences within centroid ASVs were manually added back if they increased diversity within clades (e.g. different studies, sampling locations, substrates) resulting in a final list of 37 sequences which were aligned with MAFFT using default settings (Katoh and Standley, 2013) and trimmed with trimAl (Capella-Gutiérrez et al., 2009) with a gap threshold of 0.3 and a similarity threshold of 0.001. The final sequence alignment of 1698 nucleotides was analysed with IQ-TREE (Minh et al., 2020) using ModelFinder (Kalyaanamoorthy et al., 2017) and ultrafast bootstrap with 1,000 replicates (Hoang et al., 2018). The best-fit model of substitution determined from BIC was TIM2+F+R3. The maximum likelihood tree was visualised using Treeviewer (Bianchini and SánchezBaracaldo, 2024), using Labyrinthulomycetes_LAB1 sequences as an outgroup. The identity of the host diatom ASV was determined by a manual BLAST search of the sequence against the NCBI nucleotide collection (nt). All R and shell scripts used for phylogenetic analyses are available at https://github.com/clarencesimple/SIM_Labyrinthula_merlionensis/.

### Light microscopy

A time series of the co-culture was captured for a 24 days period using Evos XL Core inverted microscope to observe changes in cell-to-cell interactions upon culture refreshment. Cell length and width were measured from 100 randomly selected vegetative cells. As *Labyrinthula* can form dense aggregates that distort cell shape and makes the cell outline unclear, only vegetative cells with distinct spindle-shaped outline were chosen for measurements. Images of the co-culture were taken using Olympus CKX53 inverted Microscope with a 40X objective and analysed using ImageJ (Schneider et al., 2012) for cell measurements.

### Scanning electron microscopy

Freshly transferred co-culture was distributed across a 24-well plate containing 8 mm poly-L-lysine coated coverslips. The plate was incubated in 22 °C 12h:12h light-dark cycles for 10 days. Four fixation methods were tested: standard protocol, standard + osmium tetroxide (OsO_4_) with glutaraldehyde (GA), standard + L-lysine and methanol-plunge. The standard protocol involved addition of a doublestrength fixative to the cultures in a 1:1 ratio; this fixative was a combination of 2x PHEM buffer with 1 M NaCl, 8% methanol-free formaldehyde, and 1.8% glutaraldehyde (GA), with the pH adjusted to growth conditions. In the case of standard + OsO_4_-GA fixation, the process was identical to the standard fixation, with the simultaneous addition of OsO_4_ to a final concentration of 0.3%. For standard + L-lysine fixation, L-lysine was added to the mix after the standard protocol to a final concentration of 75 mM. Samples treated chemically were incubated in the same 24-well plate overnight, which could result in the vapour from OsO_4_ affecting samples not intended to be treated with OsO_4_. Post-fixation, samples were rinsed in distilled water for 10 min followed by a dehydration process carried out in a series of ethanol dilutions with increased concentrations. Initially, samples were immersed in 50% solution for 20 min. This step was repeated with 70%, 80% and 90% ethanol solutions, each for 20 min, with a single repetition for every concentration. Afterwards, samples were immersed in 96% ethanol for 20 min, this time repeated twice. Finally, the samples were immersed in 100% dry ethanol for 20 min, and this step was repeated three times to ensure complete dehydration. The Methanol-plunge fixation method involved the direct plunge of samples into pure methanol for 30 sec, followed by immediate Critical Point Drying (CPD). For all fixations, CPD was performed over 8 cycles, with pressure release across two days.

### Transmission Electron Microscopy

To allow adhesion of *Labyrinthula* sp. on a flat surface for Transmission Electron Microscopy (TEM), 15 mL of a freshly transferred co-culture was incubated over a poly-L-lysine coated microscope glass slide placed in a 100 mm x 15 mm Petri dish. The sample was incubated in 22 °C 12H:12H lightdark cycles for 21 days before fixing with glutaraldehyde (1% final concentration). Using an inverted microscope, areas with high growth of *Labyrinthula* cells were marked with circles on the glass slide using a glass cutter scratch pen.

The glass slide was rinsed with cacodylate buffer for 15 mins, and fixed with 1% OsO_4_ in cacodylate buffer for 1 hour. The glass slide was then rinsed twice with cacodylate buffer, for 10 mins each, to remove residual OsO_4_. Cells on the glass slide were dehydrated in a series of increasing concentrations of ethanol (30% for 5 mins; 50% for 10 mins; 70% for 10 mins thrice; 90% for 10 mins thrice; 100% for 10 mins). Finally, 2 mL microtubes containing Spürr resin were inverted and fixed on areas on the glass slide marked with high cell concentrations. The sample was incubated at 60 °C for polymerization to take place overnight. Resin blocks on the glass slide were removed by heating the glass slide on a heating block at 50°C. Blocks were cut with an ultra-microtome (LEICA Ultracut UCT Microtome), and 60 nm-thick sections were contrasted with 2.5% uranyl acetate and 0.2% lead citrate, and observed under a JEOL 1400 TEM.

### Confocal Microscopy

Single-well glass bottom imaging chambers (Nunc LabTek II; alternatively ibidi *µ*Slides work equally well), were used to prepare the co-culture for confocal imaging. The glass chambers were first coated with 0.01% poly-L-lysine to promote cell adhesion onto the glass surface. The chambers were then rinsed with distilled water and left to dry on a heating plate at 45°C. From a newly transferred co-culture, 3 mL were transferred into each glass chamber and incubated at 22°C 12H:12H lightdark cycles. The samples were regularly observed using an inverted light microscope to follow the progression of interactions between the cells in the co-culture. The late stages of infection (Days 15 and 21), as indicated by higher concentrations of *Labyrinthula* cells, were selected for confocal imaging.

On the day of imaging, samples were fixed with 0.2 *µ*m filtered seawater (FSW) containing final concentrations of 1% para-formaldehyde (Electron Microscopy Sciences, ref. 15714) and 0.25% glutaraldehyde (Sigma, ref. G5882-100ML) for 15 mins. The fixatives were carefully removed from the glass chambers using a pipette, ensuring the cells were still adhering to the bottom surface. The samples were rinsed with FSW. Cell silica frustules were first labelled using 2 mL of 0.6% PLL-AlexaFluor54 in FSW for 15 mins in the dark, (Colin et al., 2017; Walde et al., 2023). The samples were rinsed again with FSW. A mixture of 2 mL FSW containing 0.04% DiOC3(6) and 0.6% Hoechst-33342 was used to label cellular membranes and nuclei, respectively. The samples were left to incubate in the dark for 30 mins. Finally, the samples were rinsed and kept in FSW for imaging.

A confocal microscope (Leica SP8, Germany) equipped with 40X water objective was used. At every Z-plane, fluorescence was read out in 2 sequences with 2 channels each as follows: (1a) Autofluorescence targeting Chlorophyll *a* with excitation at 405 nm and emission at 680-720 nm and (1b) DiOC3(6) targeting cellular membranes with excitation at 488 nm and emission at 505-520 nm; (2a) Hoechst-33342 targeting cell nuclei with excitation at 405 nm and emission at 415-475 nm and (2b) PLL-AlexaFluor546 targeting silica frustules with excitation at 552 nm and emission at 570-590 nm. Visualisation of 3D datasets were generated in Imaris 3D (Oxford Instruments, UK).

## RESULTS AND DISCUSSION

A *Labyrinthula* strain was isolated alongside the diatom *Biddulphia* from Singapore Straits seawater. The *Labyrinthula*-diatom co-culture (RCC7798) was subsequently established, enabling both molecular and morphological characterization of this novel Labyrinthulid, which serves as a model organism for studying infection of diatoms in tropical waters.

### Molecular characterization

#### 18S rRNA phylogeny

The full-length 18S rRNA gene sequence for *Labyrinthula* sp. from the co-culture was obtained using PacBio long read sequencing. A phylogenetic tree of the genus *Labyrinthula* was constructed using 18S rRNA sequences annotated as *Labyrinthula* spp. in the reference database PR^2^ (Guillou et al., 2012). The tree revealed distinct clades comprising sequences that correspond to specific environmental conditions or substrates (Figure 1 and Table S3). The *L. terrestris* clade comprised sequences from various studies that sampled multiple types of terrestrial grasses, including *Spartina alterniflora* (cordgrass), *Poa* spp. (bluegrass), and other turfgrasses (Table S3). The deep-branching *L. zostera* clade comprised isolates originating from multiple species of infected seagrasses (Table S3) and was previously described as a pathogenic clade linked to seagrass wasting disease (SWD) (Martin et al., 2016). All other clades are likely non-pathogenic.

**Figure 1:**
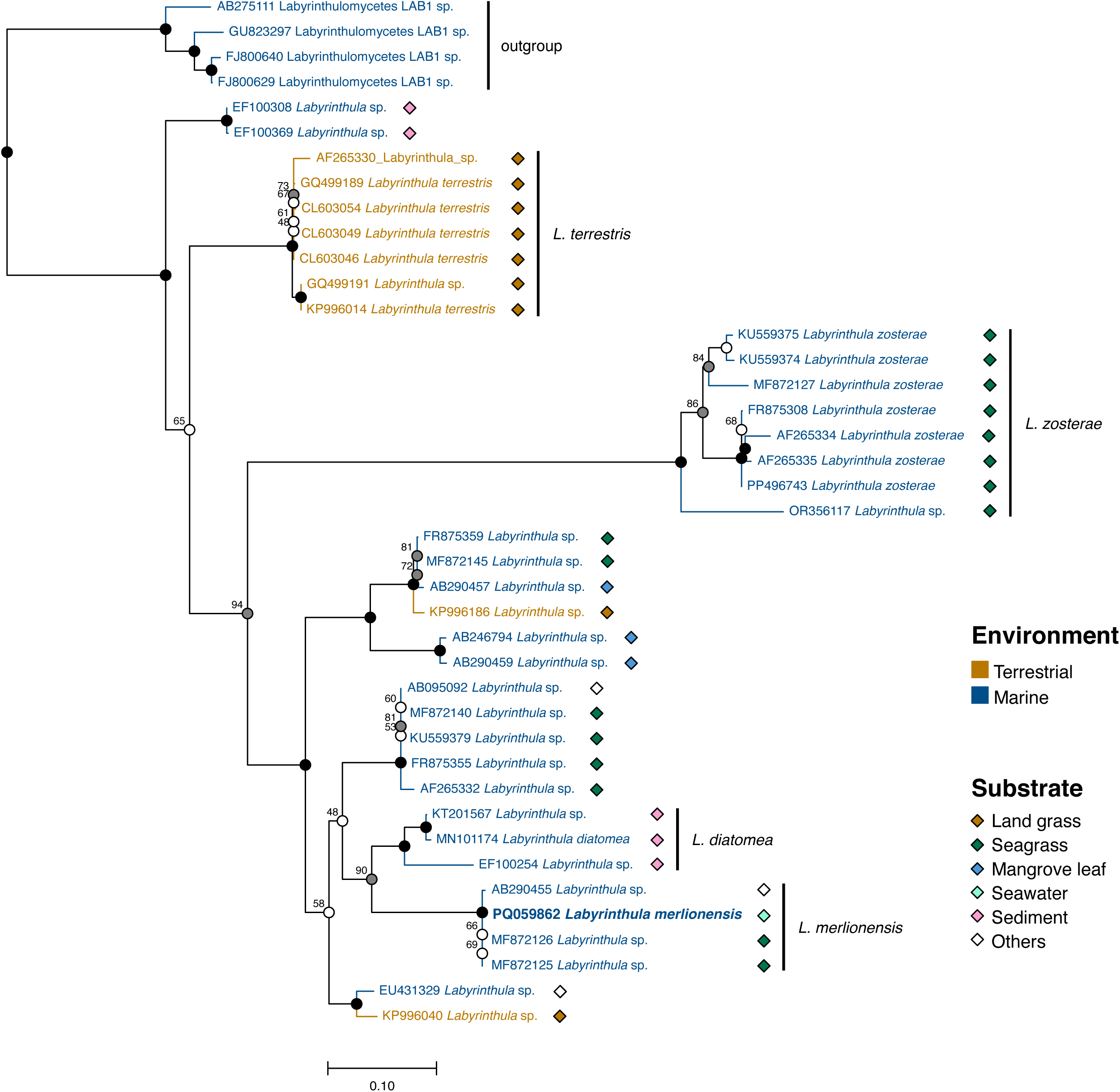
Maximum-likelihood (ML) phylogenetic tree of Labyrinthula based on clustered 18S sequences. Branches are labelled with bootstrap support. Solid dots correspond to significant support (> 95%) for ML analysis. When ML support is below 95% the percentage is indicated next to the symbol. Grey dots correspond to ML support of 70-95% Empty dot corresponds to ML support < 70%. Diamonds beside sequence names indicate the substrate where the sample was found. The strain described in this study is in bold.

Phylogenetic analysis provided strong support for our *Labyrinthula* strain (GenBank PQ059862) to belong to a clade distinct from *L. diatomea* (Figure 1). Isolates within the *L. diatomea* clade were exclusively found in sediment samples, suggesting an association with benthic diatoms and sedimentation processes (Figure 1 and Table S3). This suggests that our strain corresponds to a new species that we call *Labyrinthyla merlionensis* (see taxonomic section at the end of this part). Within its clade, *Labyrinthula* sp. (GenBank AB290454) was isolated from unidentified macroalgae, though it was cultivated on agar with the live diatom *Chaetoceros ceratosporum* (Sakata and Iwamoto, 1995; Wahid et al., 2007) (Table S3) suggesting this other *L. merlionensis* strain also relies on diatoms as a source of nutrient.

#### Diatom host identity

The full-length 18S rRNA gene sequence for the diatom host was also obtained from the co-culture (RCC7798) using PacBio (GenBank PQ059863). BLAST analysis against the NCBI nucleotide database indicated that the best matching reference cultures were CCMP147 (*Biddulphia* sp., GenBank EF585585) with 99.72% similarity (100% query coverage), and MACC6 (*Biddulphia* sp., GenBank KR007589) with 99.83% similarity (99% query coverage). The diatom was thus identified as *Biddulphia* sp. Species level characterisation of this diatom would require a combination of multigene phylogenetic analysis and detailed morphological inspection (Ashworth et al., 2013; Dąbek et al., 2017).

### Morphological characteristics of *Labyrinthula merlionensis*

Bi-flagellated zoospores (heterokont) and spindle-shaped vegetative cells that form ectoplasmic nets were commonly observed in the co-culture (Figures 2A,D). Morphological observations of *L. merlionensis* zoospores matched the distinctive characteristics of a stramenopilous heterokont (Bennett et al., 2017). The zoospore was captured in SEM only when samples were fixed using the standard protocol. The zoospore comprised a smooth, short flagellum (4 *µ*m) and a long flagellum (12 *µ*m) laterally lined with two parallel rows of fine mastigonemes (Figures 3A). The cell body did not appear to be coated by scales, which differentiates Labyrinthulida zoospores from Thraustochytrida zoospores, the latter being coated by small scales (Bennett et al., 2017). The tripartite tubular hairs at the end of each mastigoneme were not clearly captured in this micrograph, possibly due to the effects of vaporized OsO_4_ during the incubation step when preparing the samples for SEM. We ruled out the possibility that it is the diatom gamete, as centric diatom gametes are uniflagellate (Jensen et al., 2003).

**Figure 2:**
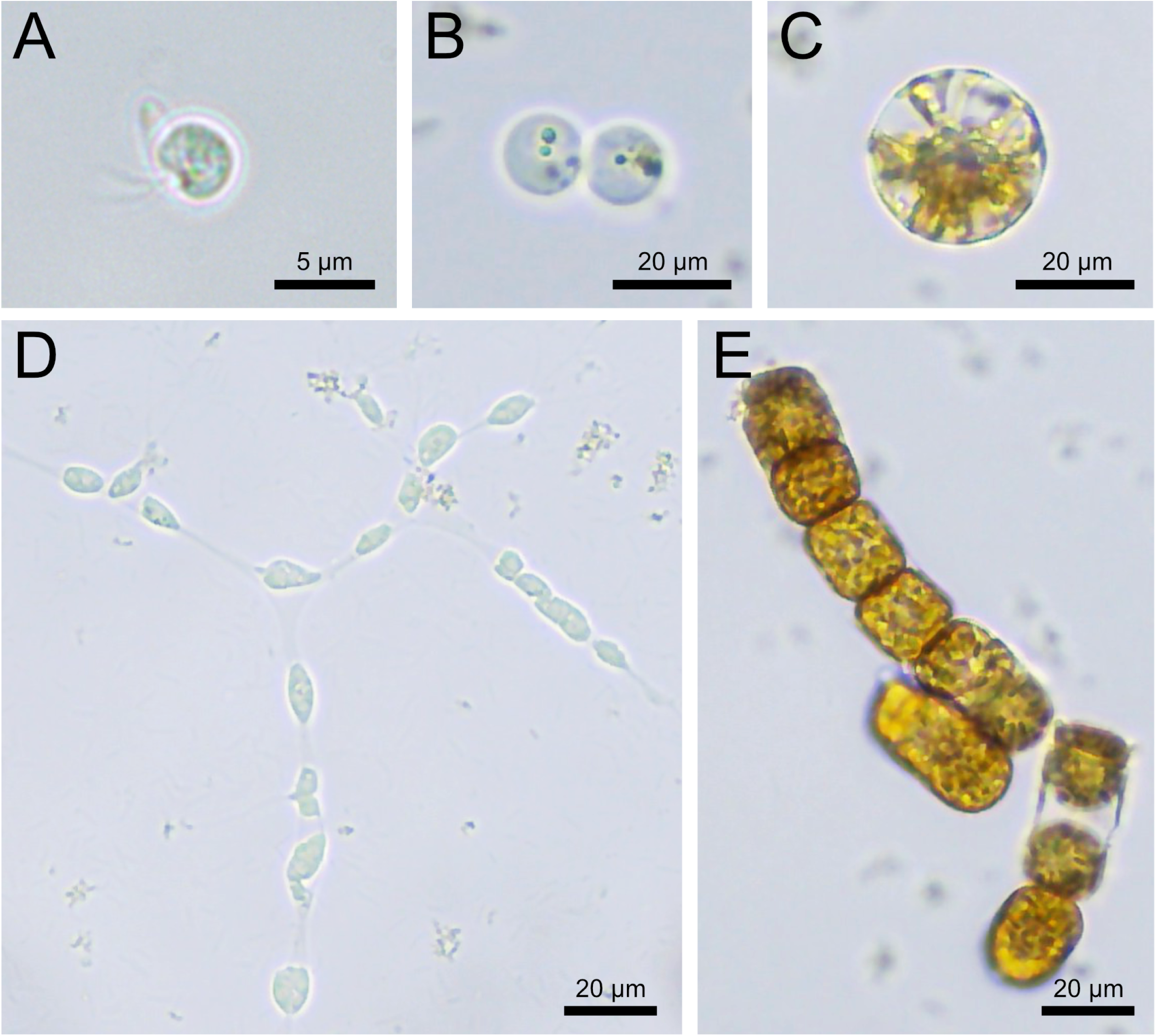
Cell types in RCC7798 co-culture. **A.** *Labyrinthula merlionensis* zoospore. **B.** Diatom spermatocytes. **C.** Diatom auxospore. **D.** Network of *L. merlionensis* vegetative cells. **E.** Two *Biddulphia* sp. chain colonies. Cell in bottom colony undergoing binary fission.

**Figure 3:**
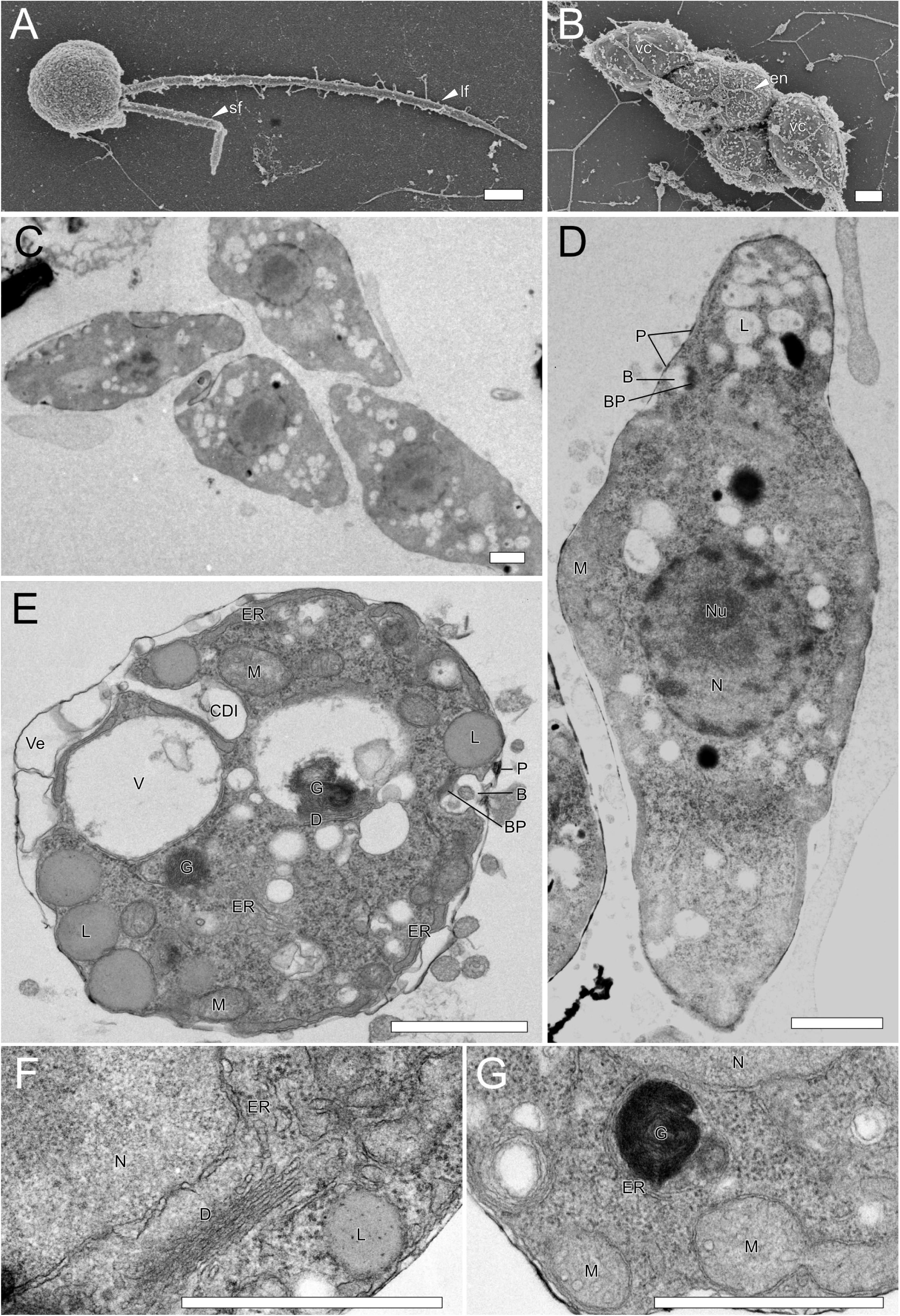
**A-B.** Scanning electron micrographs of *L. merlionensis* (RCC7798). **A.** Bi-flagellate *Labyrinthula* zoospore with a short flagellum (sf) and long flagellum (lf) lined with lateral mastigonemes. **B.** Four vegetative cells (vc) bounded by fine ectoplasmic nets (en). **C-G.** Transmission electron micrographs of thin sections of *L. merlionensis* (RCC7798). **C.** Longitudinal section of four uninucleate cells in the extracellular matrix within an ectoplasmic network. **D.** Longitudinal section of a single cell with a centrally located nucleus (N) containing the nucleolus (Nu), numerous lipid granules (L) concentrated around the nucleus and the end of the cell where the bothrosome (B) is located. A basal plaque (BP) is present at the bothrosome (B) that is adjacent to an extracellular plaque (P). **E.** Transverse section showing lipid granules and mitochondria (M) at the cell periphery, two unidentified densely coiled granular body (G), a Golgi dictyosome (D), two vacuoles (V), extracellular vesicles (Ve) and a cell division invagination (CDI). The endoplasmic reticulum (ER) mostly lies under the cell membrane. The bothrosome with its basal plaque is also present. **F.** Proximity between the nucleus, endoplasmic reticulum and Golgi dictyosome. **G.** Transverse section of an unidentified densely coiled granular body surrounded by endoplasmic reticulum and unbranched (left) and branched (right) mitochondria. Scale bars = 1 *µ*m.

The most recent published micrograph of a *Labyrinthula* zoospore was captured more than 55 years ago (Amon and Perkins, 1968) from a strain isolated from the cordgrass *Spartina alterniflora*, the same host substrate as AF265330 (Table S3) which is positioned in the *L. terrestris* clade (Figure 1). Zoospores of the SWD pathogen *L. zosterae* do not seem to have ever been observed or imaged (Sullivan et al., 2023). Zoospores of the diatom-associated *L. diatomea* were also not observed (Popova et al., 2020). Based on these records, we suggest that the formation of zoospores by *Labyrinthula* spp. could vary as a function of substrates and environmental conditions but could also be linked to genetic variation. The lack of zoospore observations in the past decades had left a significant gap in understanding the zoosporic life stage of *Labyrinthula* and its role in dispersal, settlement, and the initiation of diseases (Sullivan et al., 2023).

Vegetative *L. merlionensis* cells had width of 3-5 *µ*m and length of 6.5-10.5 *µ*m (Table 1 and Figure S2), which is smaller than the closely related *L. diatomea* (3.5–5 *µ*m width and 10.0–12.5 *µ*m length) (Popova et al., 2020). Cells within the colony, also commonly referred to as network, were connected and wrapped around by an ectoplasmic net as observed from the SEM micrograph (Figure 3B). The extracellular slime-like matrix was observed in the longitudinal section across four vegetative cells within the same network (Figure 3C).

**Table 1:**
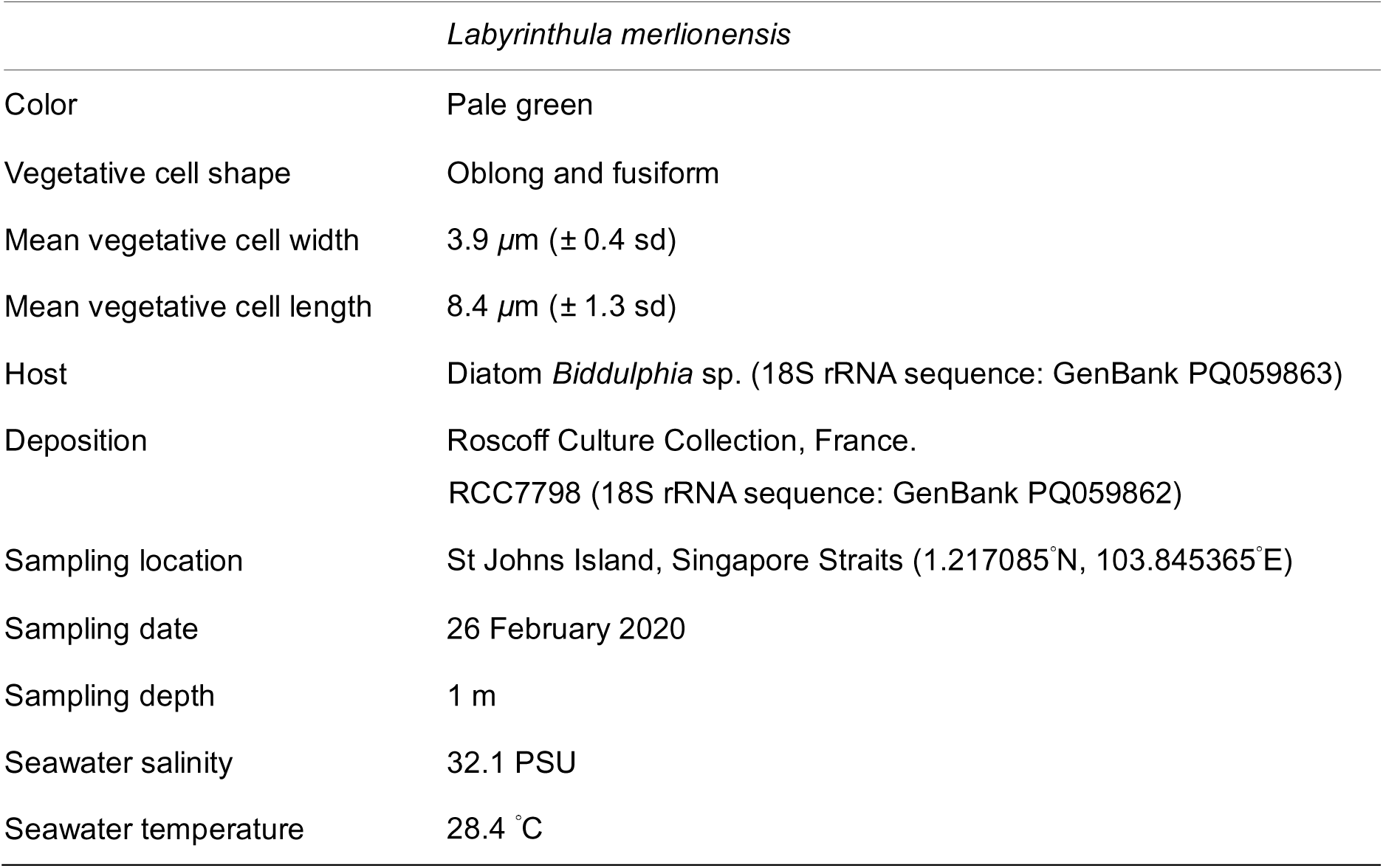
Morphological features determined from 100 vegetative cells and environmental characteristics of water sample which *Labyrinthula merlionensis* was isolated from.

The ultrastructure of *L. merlionensis* vegetative cell displayed mostly similarities with that of a *Labyrinthula* sp. strain isolated from seagrass *Zostera marina* and described by Porter (1969). The bothrosome, a cup-like invagination of the cell membrane, is a definitive characteristic of Labyrinthulida and was clearly observed in longitudinal and transverse sections of the vegetative cell plasma membrane (Figures 3D,E), with electron-dense basal plaques observed in both sections. While its exact function remains unknown, it has been hypothesized that the bothrosomes are sites of synthesis for the extracellular matrix and the network ectoplasmic membrane (Porter, 1969). The basal plaque at the bothrosome and the nearby extracellular dense plaque (Figures 3D,E) might correspond to a region possessing a high concentration of lipid, another membrane precursor (Porter, 1969). The bothrosome was also suggested to be a site of interaction between myosin from the cell body and F-actin in the extracellular matrix, resulting in their ability to glide along the ectoplasmic network (Perkins, 1973; Preston and King, 2005).

The Golgi dictyosome complex, positioned right beside the nucleus, consisted of a ribbon-shaped dictyosome made of 6 stacked and flattened cisternae (Figures 3F). In Figure 3E, the Golgi dictyosome was situated near the bothrosome, supporting the hypothesis that the Golgi is related to the formation of bothrosomes in Labyrinthulomycetes (Iwata et al., 2017). Lipid granules were not membrane-bound and were situated at the periphery of the cell (Figure 3) or around the nucleus. Branched and unbranched mitochondria were abundantly distributed around the cell (Figures 3E and G). Densely coiled unidentified granular bodies with unknown function and composition (Figures 3E and G) situated next to the nucleus, were also observed by Porter (1969). Unlike the ultrastructure of the *Labyrinthula* strain described by (Porter, 1969), no bundles of parallel microtubules were observed in our samples. This is likely due to the fact that we could not observe longitudinal sections of dividing daughter cells, as the cytoplasmic microtubules were observed to appear during cell elongation and division (Porter, 1972).

### Co-culture temporal dynamics

The development of interactions between *L. merlionensis* and *Biddulphia* was observed over 24 days following medium change of the co-culture (Figure 4). Their interactions could be characterised into three phases:

1. Establishment – Motile bi-flagellated zoospores settled onto a host or a substrate. Initial observations showed non-flagellated *L. merlionensis* zoospores physically attached to the surface of *Biddulphia* (Figure 4A) and to the glass slide (Figure 4B), indicating their ability to adhere to both biotic and abiotic surfaces. The occurrence of non-flagellated zoospores was expected, as Labyrinthulomycetes zoospores are known to lose their flagella only minutes after settlement (Iwata et al., 2017).
2. Infection – Growth of the *L. merlionensis* network and infection of diatoms. After a week, networks of vegetative cells were seen growing along the frustules of the diatom chain (Figure 4C) and on the glass slide (Figure 4D). These vegetative cells exhibited gliding movements along the ectoplasmic network, demonstrating the classic motility mechanisms of *Labyrinthula* spp. (Bennett et al., 2017). This motility is crucial for the exploration of new areas and efficient nutrient acquisition (Iwata and Honda, 2018; Porter, 1990). The increasing dominance of *L. merlionensis* resulted in more observations of dead (empty frustules) and unhealthy diatoms (weak pigmentation) (Figures 4E,F).
3. Aggregation – *L. merlionensis* network aggregated numerous diatom chains together. The *L. merlionensis* network extended their ectoplasmic nets across different diatom chains (Figures 4G,H). This included connecting unhealthy *Biddulphia* chains to healthy ones via the ectoplasmic net (Figure 4H). *L. merlionensis* did not exhibit any observable preference to colonize live, unhealthy, or dead diatom cells. A quantitative approach to follow the parasite-host dynamics would be necessary to test the “healthy herd” hypothesis, where the parasite performs selective parasitism, weeding out unhealthy diatoms to maintain the overall population health (Laundon et al., 2021). After three weeks, a dense network of vegetative cells was formed, concentrated around a large agglomerate of live and dead *Biddulphia* cells (Figure 4I).

**Figure 4:**
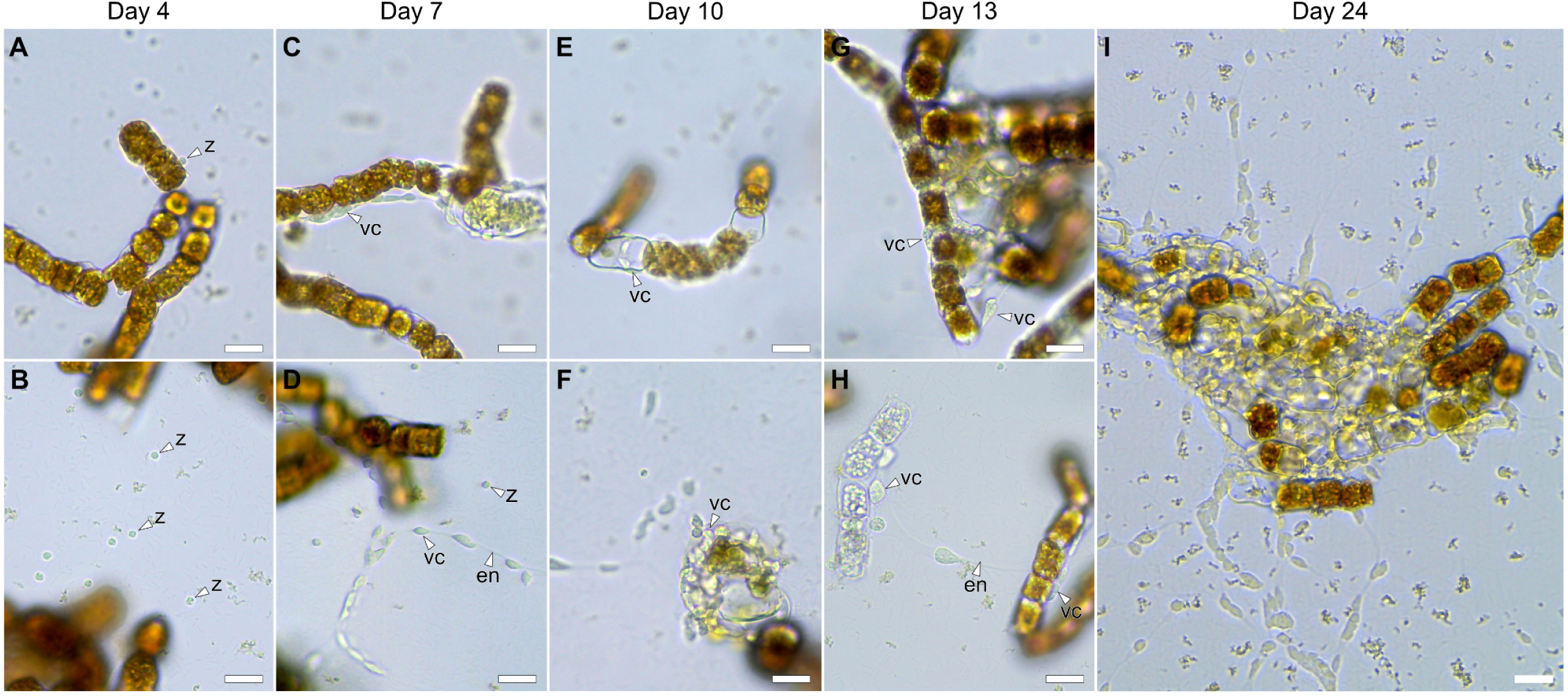
Progression of the RCC7798 *Labyrinthula merlionensis*-diatom co-culture from day of culture medium change. **A.** *L. merlionensis* zoospore physically attached to diatom. **B.** *L. merlionensis* zoospores attached to glass slide (white arrowheads). **C.** *Labyrinthula* network of vegetative cells growing along the exterior of a diatom chain. **D.** *L. merlionensis* vegetative cells gliding along the ectoplasmic network. **E.** Short network of *Labyrinthula* vegetative cells growing around an empty diatom frustule on a diatom chain. **F.** Network of *Labyrinthula* vegetative cells attached to frustules of discolored diatoms. **G.** Large network of *Labyrinthula* vegetative cells attached around an agglomeration of empty and live *Biddulphia* cells. **H.** An unhealthy *Biddulphia* colony connected physically to a healthy diatom colony via the *L. merlionensis* ectoplasmic net. **I.** A large network of dense *L. merlionensis* vegetative cells concentrated around a large agglomeration of live and dead *Biddulphia* cells. Scale bars = 20 *µ*m; z = zoospores; vc = vegetative cell; en = ectoplasmic net.

### Cell-to-cell interactions

SEM and confocal microscopy were carried out on the co-culture to better characterize cell-cell interactions during the parasitism phase. SEM observations showed the attachment of *L. merlionensis* vegetative cells on diatom chains at the marginal ridge region between diatoms (Figures 5A,C). A closer examination revealed that the ectoplasmic nets extended into the ridge and also around diatom frustules (Figures 5B,D). *L. merlionensis* might have a preference in targeting this region that could be the most vulnerable part of the diatom chain, as the diatom frustules are rigid and likely impenetrable. This targeting behaviour is consistent with findings in another Labyrinthulida, *Aplanochytrium*, which adheres to living diatom cells via the tips of its ectoplasmic net system to extract and absorb nutrients, rather than releasing digestive enzymes like saprophytes (Hamamoto and Honda, 2019). This contrasts with another order of Labyrinthulomycetes, the Thraustochytrida (e.g. *Schizochytrium aggregatum*), which secrete the hydrolytic enzyme carboxymethyl cellulase via the bothrosome to digest the host’s organic components, allowing for the penetration of ectoplasmic nets for direct intake of glucose (Iwata and Honda, 2018; Perkins, 1973).

**Figure 5:**
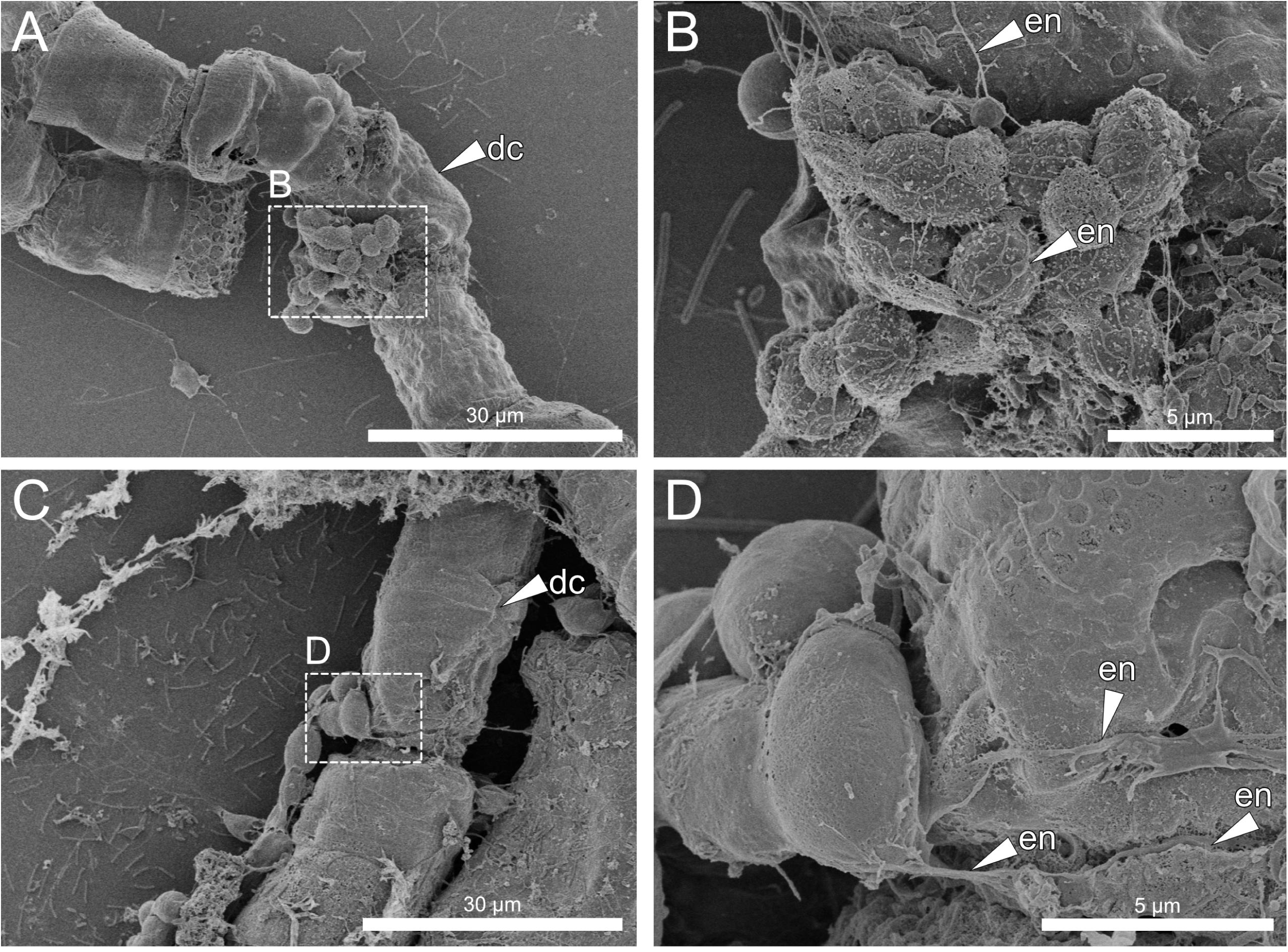
Scanning electron micrographs of *Labyrinthula merlionensis* (RCC7798). **A.** Aggregate of *L. merlionensis* cells on a diatom chain. **B.** *L. merlionensis* cells adhering to each other and on the diatom chain with fine ectoplasmic nets. **C.** A long network of *L. merlionensis* cells attached to a diatom chain. **D.** Fine *L. merlionensis* ectoplasmic net extending deep into the marginal ridge between two diatom cells and around the diatom frustule above. dc = diatom chain; en = ectoplasmic net.

The preference for colonizing the marginal ridge regions of diatom chains was also observed using confocal microscopy during the infection phase (day 15, Figures 6, 7). The presence of intact and well-defined chloroplasts inside the frustules of individual *Biddulphia* cells were used as a proxy for cell health (Walde et al., 2023). The diatom chain in Figures 6A,B comprised *Biddulphia* cells of varying health. At the marginal ridge between two healthy diatom cells, *L. merlionensis* cells were concentrated at one spot (Figure 6C). A cross-section of this marginal ridge showed the absence of nuclei or membrane signals from *L. merlionensis* within the frustule (Figure 6D). The diatom chain also comprised a dead cell (empty frustule with no chloroplasts) wrapped around by the *L. merlionsis* network (Figures 6A,B). A cross-section of the empty frustule (Figure 6E) showed the absence of nuclei or membrane signals from *L. merlionensis* inside the frustule. This indicates that there was no physical intrusion into the diatom frustule regardless whether the diatom was live or dead, supporting that *L. merlionensis* utilized its ectoplasmic nets for motility and nutrient absorption but not for penetration of host cells (Hamamoto and Honda, 2019; Iwata and Honda, 2018).

**Figure 6:**
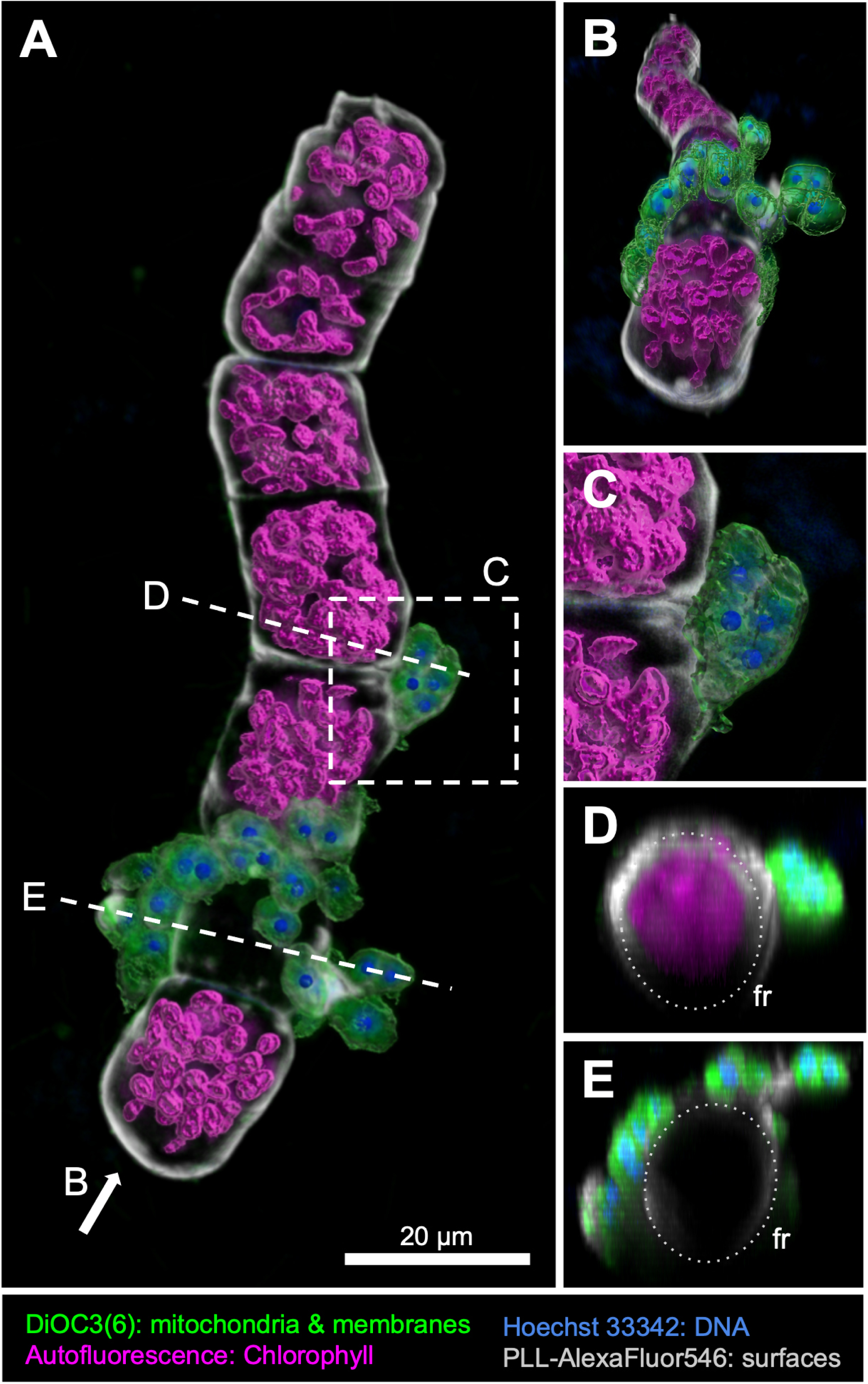
3D renderings of a confocal dataset of a 15 days old RCC7798 *Labyrinthula*-diatom co-culture focusing on one diatom chain. **A.** Network of *Labyrinthula* vegetative cells extending from the glass slide to multiple diatom cells. **B.** Frontal 3D-perspective showing the collar of vegetative *Labyrinthula* cells around the colonised diatom chain. **C.** Zoomed view of the marginal ridge between two diatom cells, with multiple vegetative *Labyrinthula* cells closely packed. **D.** Cross-section from A (dashed area seen from perspective B), showing only chlorophyll signals from within the diatom frustule (dotted circle). **E.** Cross-section from A (dashed area seen from perspective B) through an empty diatom frustule (fr; dotted circle), showing no DNA or chlorophyll signals inside the frustule.

We investigated another more intensely infected and longer diatom chain from another replicate sample and took cross-section slices of the 3D model at multiple sites of infection (Figure 7). The *Biddulphia* chain was composed of mostly healthy cells, with chloroplasts occupying most of the cell volume (Figure 7B). 3D reconstruction of the empty diatom frustule in the middle of the chain showed *L. merlionensis* vegetative cells and isolated nuclei within the frustule (Figure 7), suggesting a successful invasion into the empty cell. However, it was unclear if the frustule had an opening or whether any entry points were caused by penetration by ectoplasmic nets. Three other marginal ridge regions between diatom cells were also imaged (Figure 7C-E). None of the cross-sections showed any sign of intrusion by *L. merlionensis* which were only detected on the diatom external surface.

**Figure 7:**
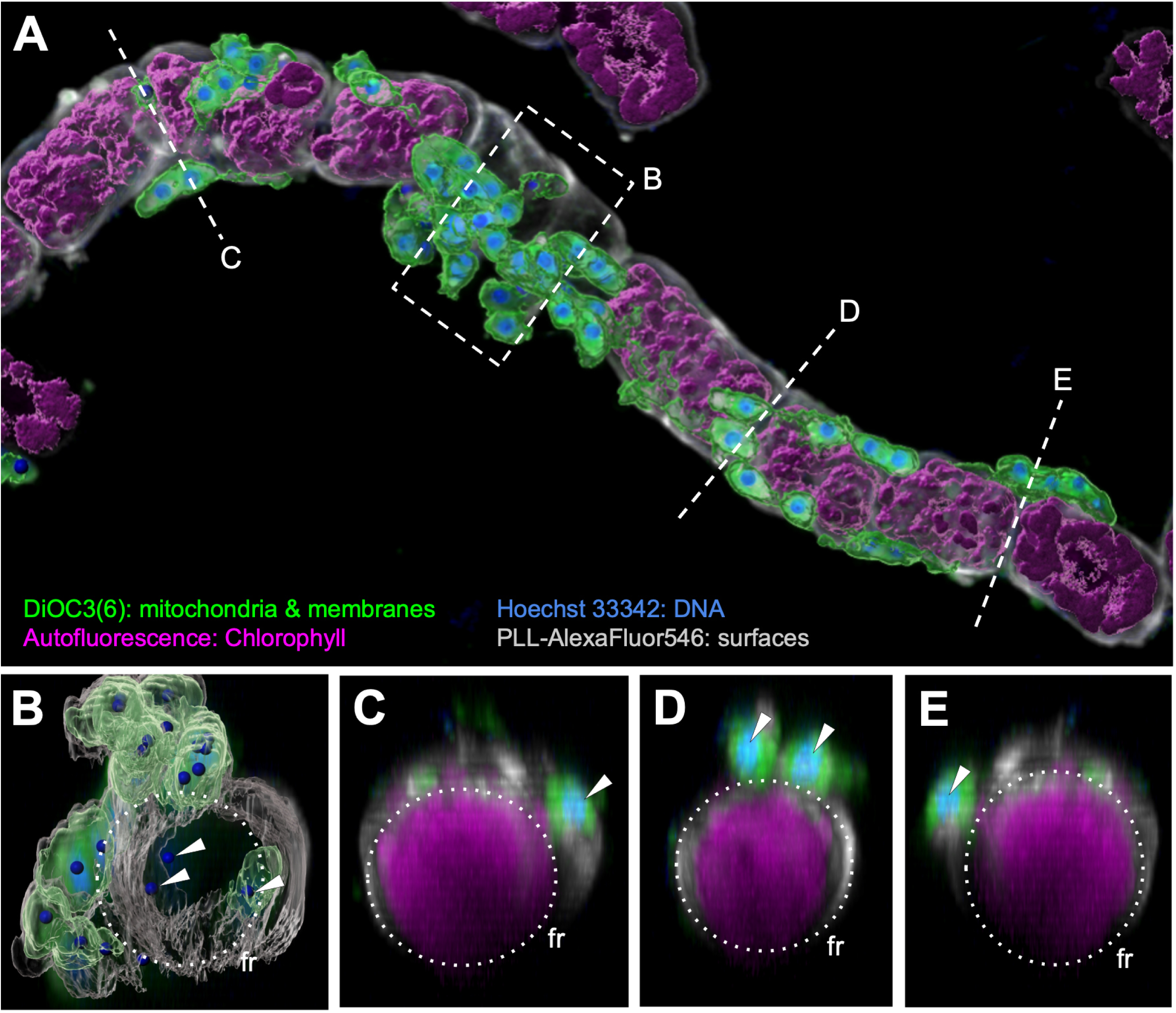
3D renderings of a confocal dataset of an infected long diatom chain from a 15 days old RCC7798 *Labyrinthula*-diatom co-culture. **A.** Extensive network of *Labyrinthula* vegetative cells on the surface of the diatom chain. **B.** Cross-section view through an empty frustule (region. Reconstructed 3D projection showed multiple strong DNA (white arrowheads) and membrane signals within the empty frustule (dotted circle). **C-E**. Cross-sections of three investigated sites of marginal ridges between two diatom cells displaying strong autofluorescence signals from chlorophyll-containing plastids (magenta) inside live diatom cells, with DNA and membrane signals (white arrowheads) of the ectoplasmic net on the exterior surface of the diatom frustule (fr; dotted circle).

Confocal microscopy of an older culture (day 21) during a later infection stage showed strong DNA signals within the frustule of unhealthy diatoms (Figure 8). The three infected diatom cells contained chloroplasts less densely packed compared to the diatoms in the longer chain below (Figure 8A). These spherical DNA signals in the infected diatom cell were the same size as *L. merlionensis* nuclei (Figures 8B-D). Since there were at least six DNA signals in the infected diatom (Figures 8B-D), nucleic DNA signals from diatom auxospore were ruled out. if it were the case, there would have been a maximum of three nuclei — one functional nucleus and one or two pyknotic nuclei, that are derived from the acytokinetic mitosis preceding the formation of each valve (Kaczmarska et al., 2022). No membrane signals in the shape of spindle-shaped vegetative *L. merlionensis* cells were detected (Figure 8) in contrast to what is observed in Figure 7. If these nuclei signals were indeed from *L. merlionensis* zoospores, it would suggest that this strain could also be an intracellular parasite (Rueckert et al., 2019), possibly by producing zoospores in host cells.

**Figure 8:**
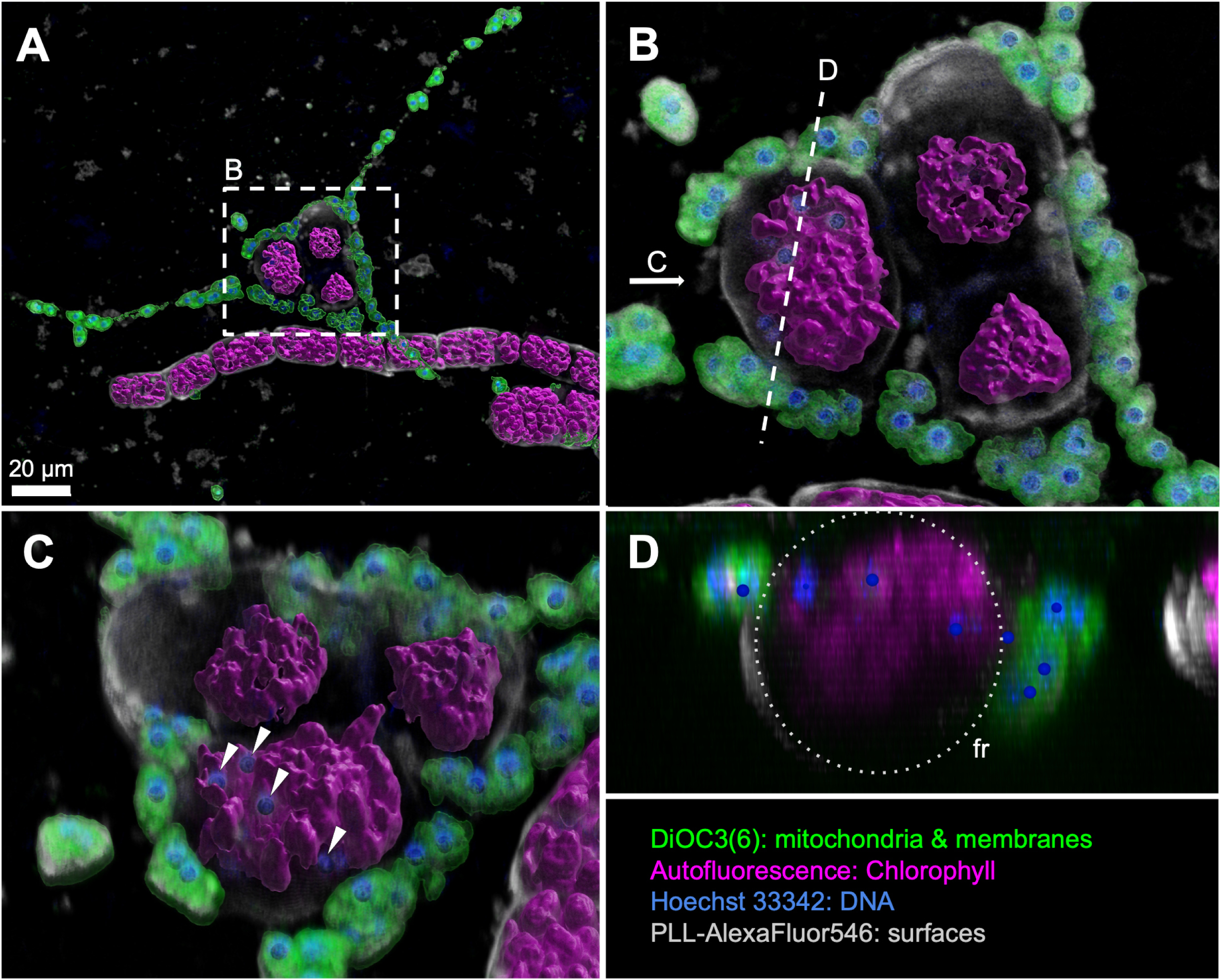
3D renderings of a confocal dataset of three infected diatom cells from a 21 days old RCC7798 *Labyrinthula*-diatom co-culture. **A.** Network of *Labyrinthula* vegetative cells extending from the glass slide to multiple diatom cells. **B.** Close-up of the three diatom cells lined with a network of vegetative *Labyrinthula* cells. **C.** Multiple strong signals of small nuclei amongst chlorophyll-containing plastids (magenta) inside a live diatom cell, indicated by blue spheres (white arrowheads). **D.** Cross-section from B (dashed area seen from the left side), showing DNA signals from within the diatom frustule (fr, dotted circle).

### Host range

Within the *L. merlionensis* clade, MF872126 and MF872125 were isolated from the surfaces of seagrasses and AB290455 was isolated from seaweed (Table S3). They were not pathogenic strains linked to SWD (Martin et al., 2016). Our strain is the only one isolated from seawater having a diatom host (Table S3). The association of strains from the *L. merlionensis* clade with microalgae (diatoms), macroalgae (seaweed) and seagrasses suggest a wide range of substrates. However, based on observed behaviour and interactions with diatoms (Figures 4-8), we speculate that *L. merlionensis* could also be found on surfaces of macroalgae and seagrasses in association with epiphytic diatoms.

The parasitic-like interaction between *L. merlionensis* and diatoms might not be restricted to the genus *Biddulphia*. The *L. diatomea* strain in the adjacent clade (Figure 1) was isolated with the diatom *Cylindrotheca closterium* and then cultivated with another diatom *Micropodiscus weissflogii* (Popova et al., 2020), demonstrating variability in host taxa. The range of host specificity for *L. merlionensis* will require further investigation by pairing the isolated *L. merlionensis* strain with various diatom taxa. Our attempts to isolate *L. merlionensis* in pure culture have been unsuccessful due to fungal contamination. However, we noted that *L. merlionensis* managed to thrive on carbonrich RS media agar plate (Sigona and Richter, 2023) prior to fungal contamination. Based on the dependence on media with carbon sources and the association of other Labyrinthulida with several species of diatoms (Hamamoto and Honda, 2019; Popova et al., 2020), we speculate that *Labyrinthula* species depend on a unique set of carbon sources that the hosts can produce.

## Conclusion

The ability to form large and dense networks (> 100 *µ*m) around both live and dead cells suggests that *L. merlionensis* plays a role in nutrient cycling (Raghukumar, 2017) and facilitates vertical carbon export (Bai et al., 2021). *L. merlionensis* can be characterized as an ectoparasite that colonizes the surface of diatoms. The wrapping of vegetative cells and their ectoplasmic nets around diatom frustules is likely to hinder growth and limits nutrient exchange between the environment and diatom cells, while outcompeting the diatoms for nutrients. We also raised the possibility that *L. merlionensis* could act as an endoparasite that produces zoospores inside the host cell. Diatoms are a very important phytoplankton group and therefore understanding the infection mechanisms of *Labyrinthula* species on diatoms can shed light on complex interactions within marine ecosystems (Bjorbækmo et al., 2020), potentially leading to insights into how these interactions affect nutrient cycling and the biological carbon pump (Bai et al., 2021; Raghukumar, 2017; Raghukumar and Damare, 2011). Incorporating controlled physiological experiments in future work, such as testing the effects of temperature, light, and pH on infection rates and modes, will help elucidate the environmental drivers and constraints shaping *Labyrinthula*–diatom interactions and provide deeper insight into the processes underlying infection dynamics.

## TAXONOMY SECTION

*Labyrinthula merlionensis* Sim and Lopes dos Santos sp. nov.

Spindle-shaped vegetative cells measure 6.5-10.5 *µ*m in length and 3-5 *µ*m in width. Unbranched and branched mitochondria and endoplasmic reticulum populate the cell periphery. A densely coiled unidentified granular body lies adjacent to the nucleus. Cells are associated with a diatom from the genus *Biddulphia*.

Type locality: Strain RCC7798 was isolated from seawater sampled in the Singapore Straits, off the west coast of Saint John’s Island (1.217085°N, 103.845365°E)

GenBank accession number: PQ059862 represents the 18S rRNA gene sequence

Holotype: Cells embedded in resin blocks were deposited at the Lee Kong Chian Natural History Museum, National University of Singapore, accession number ZRC.MIS.0074 and ZRC.MIS.0075. Authentic culture was deposited as RCC7798 at the Roscoff Culture Collection, France.

Etymology: Named after the official mascot of Singapore, where the strain was isolated. It is a mythical creature that is a physical fusion of a lion head and a scaled fish as the lower body.

## AUTHOR CONTRIBUTIONS STATEMENT

CWHS and ALS conceived the study. ALS and AK isolated and cultivated the co-culture. CWHS obtained and processed the PacBio sequences. CWHS and DV analysed the PR^2^ sequences and performed phylogenetic analyses. CWHS executed the light microscopy time series. MW and CWHS obtained and analysed the samples for confocal microscopy. SLP, PG and CWHS obtained and analysed the samples for TEM. HS and JJ obtained and analysed the samples for SEM. CWHS and ALS drafted the manuscript. All authors discussed data, read and approved the final manuscript.

## DATA AVAILABILITY

Nucleotide sequences for RCC7798 were deposited to GenBank under accession numbers PQ059862 and PQ059863. All codes and data used in this study can be found in https://github.com/clarencesimple/SIM_Labyrinthula_merlionensis/tree/main.

