## Supplementary Materials for "*Labyrinthula merlionensis* sp. nov.: a novel labyrinthulid infecting marine diatoms"

### SUPPLEMENTARY INFORMATION

**Table S1:** RCC7798 flask variations were used for DNA extraction and PacBio Sequel II sequencing, with the number of reads documented at each stage of the DADA2 processing pipeline.

| Sample name | Age of culture | DNA conc (ng $\mu L^{-1}$) | CCS reads | After filterAndTrim | denoise | After chimera removal |
|---|---|---|---|---|---|---|
| LABY01 | 84 days | 24.9 | 7194 | 5469 | 5435 | 5419 |
| LABY02 | 84 days | 30.1 | 8496 | 6511 | 6418 | 6418 |
| LABY03 | 5 days | 33.4 | 23593 | 17737 | 17571 | 17325 |
| LABY04 | 11 days | 45.6 | 6180 | 4823 | 4740 | 4703 |

**Table S2:** Taxa retrieved from PacBio Sequel II sequencing of RCC7798

| Class | Species | n CCS reads | n ASV | n clusterASV |
|---|---|---|---|---|
| Labyrinthulomycetes | *Labyrinthula* sp. | 33557 | 43 | 3 |
| Bacillariophyta | *Biddulphia* sp. | 205 | 1 | 1 |
| Cryptophyceae | *Goniomonas amphinema* | 159 | 2 | 2 |

**Table S3:** Metadata of sequences used to construct *Labyrinthula* phylogeny in Figure ??, sorted according to their position on the tree. GB = GenBank. ID code represents any form of unique identification to a. culture, isolation, or environmental sequence. Cluster size indicates the number of unique sequences clustered with this sequence at 99% which were removed to avoid redundancy. An NA value indicates they were manually added to increase strain diversity within clade. In references, DS indicates direct submission of sequence to NCBI without an accompanying article.

| | Species | GB Accession | ID code | Cluster size | Environment | Substrate | Substrate description | Location | Reference |
|---|---|---|---|---|---|---|---|---|---|
| 1 | *Labyrinthula* sp. | EF100308 | D3P06A09 | 2 | marine | sediment | Oxygen-depleted intertidal | Greenland, Disko Island, Unqussivik | Stoeck et al. (2007) |
| 2 | *Labyrinthula* sp. | EF100369 | D5P10B11 | NA | marine | sediment | Oxygen-depleted intertidal | Greenland, Disko Island, Unqussivik | Stoeck et al. (2007) |
| 3 | *Labyrinthula* sp. | AF265330 | f | 1 | terrestrial | land grass | *Spartina alterniflora* | USA, GA, Sapelo Island | Leander and Porter (2001) |
| 4 | *L. terrestris* | GQ499189 | Laby10 | 8 | terrestrial | land grass | *Poa trivialis* | USA, South Carolina | Douhan et al. (2009) |
| 5 | *L. terrestris* | CL603054 | UT1-4 | 1 | terrestrial | land grass | *Poa. annua* | USA, Utah | Craven et al. (2005) |
| 6 | *L. terrestris* | CL603049 | CA-5 | 1 | terrestrial | land grass | *P. annua* | USA, CA | Craven et al. (2005) |
| 7 | *L. terrestris* | CL603046 | AZ-3 | 6 | terrestrial | land grass | *P. annua* | USA, AZ | Craven et al. (2005) |
| 8 | *Labyrinthula* sp. | GQ499191 | NA | 7 | terrestrial | land grass | *P. trivialis* | USA, AZ, Tucson, golf course | Douhan et al. (2009) |
| 9 | *L. terrestris* | KP996014 | Laby830 | NA | terrestrial | land grass | Turfgrass | USA, AZ and New Mexico | Chitrampalam et al. (2015) |
| 10 | *L. zosterae* | KU559375 | 302b9-PO | 1 | marine | seagrass | *Posidonia oceanica* | Spain, Alicante, Alicante Bay | Martin et al. (2016) |
| 11 | *L. zosterae* | KU559374 | 18b-CN | 1 | marine | seagrass | *Cymodocea nodosa* | USA, VA, Chesapeake Bay, Cape Charles | Martin et al. (2016) |
| 12 | *L. zosterae* | MF872127 | DP1_Zm_F | 4 | marine | seagrass | *Zostera muelleri* | Spain, Mallorca Island, Pollenca | Trevathan-Tackett et al. (2018) |
| 13 | *L. zosterae* | FR875308 | LA79 | 22 | marine | seagrass | *Zostera marina* | Italy, Adriatic Sea, Gabicce Mare | Bockelmann et al. (2012) |
| 14 | *L. zosterae* | AF265334 | MBL_93-2 | 1 | marine | seagrass | *Z. marina* | USA, MA, Woods Hole | Leander and Porter (2001) |
| 15 | *L. zosterae* | AF265335 | type-154 | 1 | marine | seagrass | *Z. marina* | USA, WA, San Juan Island | Leander and Porter (2001) |
| 16 | *L. zosterae* | PP496743 | V24 | NA | marine | seagrass | Z. marina | USA, Oregon | Agnew et al. (2024) DS |
| 17 | *Labyrinthula* sp. | OR356117 | L7.2 | 1 | marine | seagrass | *Thalassia testudinum* | USA, Florida | Ugarelli et al. (2024) |
| 18 | *Labyrinthula* sp. | FR875359 | Sva13 | 8 | marine | seagrass | *Z. marina* | Finland, Northern Baltic, Svartholm | Bockelmann et al. (2012) |
| 19 | *Labyrinthula* sp. | MF872145 | Ln5 | NA | marine | seagrass | *P. australis* | Australia, New South Wales, Palm Beach | Trevathan-Tackett et al. (2018) |
| 20 | *Labyrinthula* sp. | AB290457 | 01-Jy-1b | NA | marine | others | Mangrove leaf | Japan, Kagoshima Bay | Wahid et al. (2007) |
| 21 | *Labyrinthula* sp. | KP996186 | Laby2020 | 1 | terrestrial | land grass | Turfgrass | USA, AZ and New Mexico | Chitrampalam et al. (2015) |
| 22 | *Labyrinthula* sp. | AB246794 | N8 | 10 | marine | others | Mangrove leaf | Japan | Tsui et al. (2009) |
| 23 | *Labyrinthula* sp. | AB290459 | 00-Bat-05 | 1 | marine | others | Mangrove leaf | Philippines, Batan Bay | Wahid et al. (2007) |
| 24 | *Labyrinthula* sp. | AB095092 | L59 | 9 | marine | others | Unidentified floating leaf | Japan, Hokkaido | Kumon et al. (2003) |
| 25 | *Labyrinthula* sp. | MF872140 | Ln11 | NA | marine | seagrass | *Z. nigricaulis* | Australia, Victoria, Portarlington | Trevathan-Tackett et al. (2018) |
| 26 | *Labyrinthula* sp. | KU559379 | 215b-ZM | | marine | seagrass | *Z. marina* | USA, VA, Chesapeake Bay, Cape Charles | Martin et al. (2016) |
| 27 | *Labyrinthula* sp. | FR875355 | San25 | | marine | seagrass | *Z. marina* | Norway, Skagerrak, Sandspollen | Bockelmann et al. (2012) |
| 28 | *Labyrinthula* sp. | AF265332 | s | 1 | marine | seagrass | *Z. marina* | USA, NH, Adams Pt. | Leander and Porter (2001) |
| 29 | *Labyrinthula* sp. | KT201567 | QZ.18S 5 | 5 | marine | sediment | Benthic diatom film | Unknown | Huang & Shao (2015) DS |
| 30 | *L. diatomea* | MN101174 | NA | NA | marine | sediment | Sediment sample | Thailand, Pattaya | Popova et al. (2020) |
| 31 | *Labyrinthula* sp. | EF100254 | D2P04F11 | 1 | marine | sediment | Oxygen-depleted intertidal | Greenland, Disko Isalnd, Unqussivik | Stoeck et al. (2007) |
| 32 | *Labyrinthula* sp. | AB290455 | L95-1 | 6 | marine | others | Seaweed | Japan, Kagoshima Bay | Wahid et al. (2007) |
| 33 | ***L. merlionensis*** | **PQ059862** | RCC7798 | NA | marine | seawater | Cultivated with *Biddulphia* | Singapore, Singapore Straits | This study |
| 34 | *Labyrinthula* sp. | MF872126 | WAR Zm A | NA | marine | seagrass | *Z. muelleri* | Australia, Victoria, Warneet | Trevathan-Tackett et al. (2018) |
| 35 | *Labyrinthula* sp. | MF872125 | DP1_Pa_G | NA | marine | seagrass | *P. australis* | Australia, Victoria, Duck Point | Trevathan-Tackett et al. (2018) |
| 36 | *Labyrinthula* sp. | EU431329 | LTH | 2 | marine | others | Amoeba symbiont in fishgill | Spain, Northwest farm | Dyková et al. (2008) |
| 37 | *Labyrinthula* sp. | KP996040 | Laby879 | 1 | terrestrial | land grass | Turfgrass | USA, AZ and New Mexico | Chitrampalam et al. (2015) |

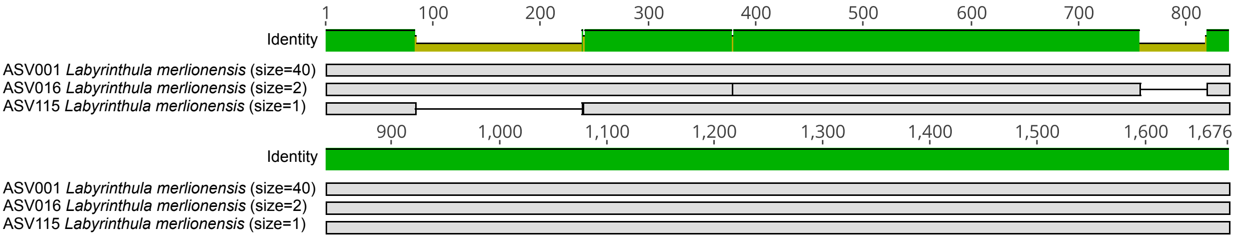


**Figure S1:** Alignment of three centroid ASVs of *L. merlionensis* sp. 18S rRNA gene. Size indicates number of ASVs clustered with the centroid sequence based on 99% similarity.

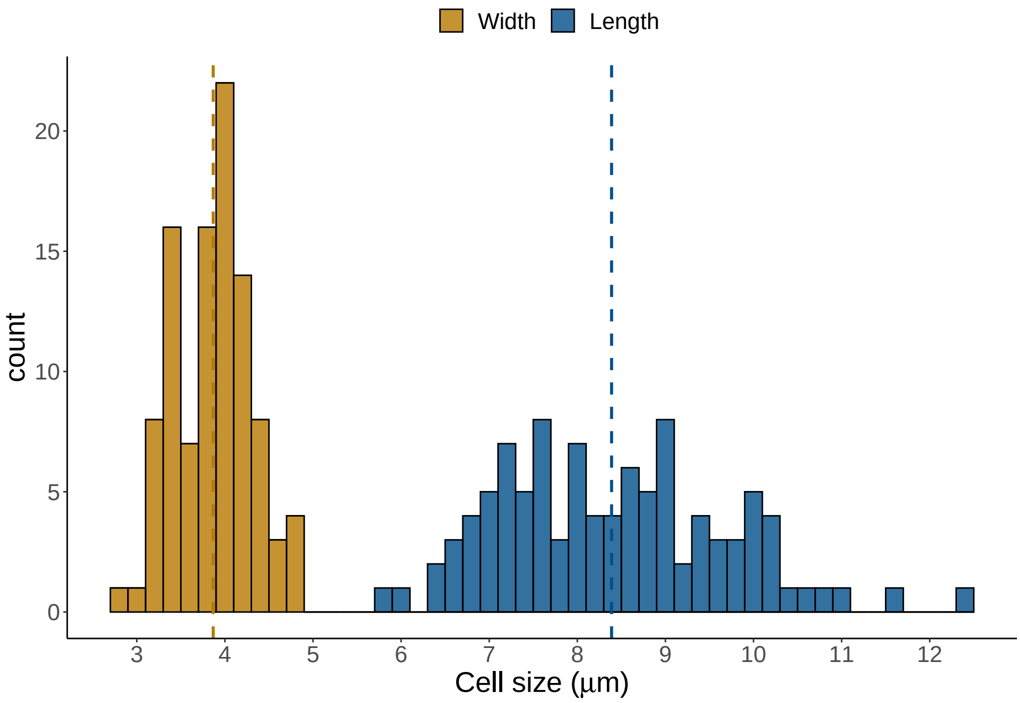


**Figure S2:** Cell size based of 100 vegetative *L. merlionensis* cells. Histogram counts were binned at 0.2 $\mu$m. Dashed line indicates means of cell width and length.
